# Capture and Purification of Plant Genomic DNA on a Readily-available Cellulose Matrix

**DOI:** 10.1101/163212

**Authors:** Megan M. Thompson, Estelle M. Hrabak

**Affiliations:** Department of Molecular, Cellular and Biomedical Sciences University of New Hampshire 46 College Road Durham, NH 03824

**Keywords:** FTA® Cards, PCR, plant genomic DNA purification, genotyping

## Abstract

Whatman FTA ®Cards are a fast and efficient method for capturing and storing nucleic acids but are cost-prohibitive for some researchers. We developed a method that substitutes readily-available cellulose-based paper and homemade washing buffer for commercial FTA ®Cards and FTA ®Purification Reagent. This method is suitable for long-term storage of DNA from many plant species, including *Arabidopsis thaliana*, prior to purification and PCR.

**Method Summary:** Here we report a low-cost method for long-term storage of plant genomic DNA on a readily available cellulose matrix.

---

In protocols where large numbers of DNA samples are collected, stored, and analyzed by PCR, decreasing the time and/or cost associated with preparing the DNA is desirable. For *Arabidopsis thaliana*, the method of Klimyuk, et al. (1) is quick and low-cost but, in our hands, samples usually become less reliable DNA templates after several weeks of storage at 4 ° C or -20 ° C. Whatman FTA^®^ Cards (GE Healthcare Life Sciences, Pittsburgh, PA) are often used to collect and store biological samples on paper impregnated with proprietary chemicals (2-5). Samples such as body fluids or plant tissue are pipetted or imprinted onto the FTA^®^ Cards, then dried completely for storage. When DNA is needed for PCR, small disks are removed from the card and the disks are washed twice with commercial FTA^®^ Purification Reagent and then twice with TE_0.1_ (10 mM Tris-Cl, pH 8.0, 0.1 mM EDTA). Disks are dried completely at room temperature for one hour before adding PCR reagents. While this method is relatively fast and efficient, the FTA^®^ Purification Reagent and FTA^®^ Cards are costly. We investigated reliable and cost-effective substitutions for these materials.

In place of 200 μL FTA^®^ Purification Reagent to wash the FTA^®^ paper disks, we tested the same volume of five different solutions: homemade TENT buffer (10 mM Tris-Cl, pH 8.0, 1 mM EDTA, 12 mM NaCl, and 2.5% Triton X-100), 20 mM NaOH, 1% SDS, sterilized distilled water, or TE_0.1_. Both 20 mM NaOH and 1% SDS were previously reported as alternative washing solutions (6). PCR was used to amplify a 650 bp product from an Arabidopsis protein phosphatase gene.Disks that were washed with water, TE_0.1_, NaOH or SDS yielded inconsistent results (data not shown). Consistent product was generated from disks washed with either TENT buffer or commercial FTA^®^ Purification Reagent (data not shown); thus, TENT buffer was used for the initial two washes in all subsequent experiments.

Next, we investigated alternate matrices and final wash solutions. DNA tightly associates with cellulose fibers (7), so we investigated three cellulose-based matrices as alternatives to commercial FTA^®^ Cards: untreated Grade 1 filter paper (Whatman) and either untreated or untreated grade 238 chromatography paper (VWR International, Radnor, PA). Treated paper was prepared by soaking in 400 mM Tris-Cl (pH 8.0), 5% sodium dodecyl sulfate (SDS), and 25 mM EDTA for 2 h, followed by drying at room temperature overnight. Treated paper can be stored for at least 6 months prior to use. Optimization of the method was performed using leaves from 4 week-old *Arabidopsis thaliana* ecotype Columbia-0. The cellulose matrix was sandwiched between pieces of parchment paper to prevent contamination of tools during tissue printing and to avoid physical contact between samples during storage. The plant tissue was pressed into the paper using moderate pressure from a ceramic pestle. Leaf imprints on the cellulose matrix were dried for at least one hour before punching a 1.5-mm disk using a Miltex biopsy punch with plunger (Ted Pella, Inc., Redding, CA). We tested several alternatives to TE0.1 for the final two washes: sterilized deionized water, 100% ethanol, or 100% isopropanol. After the final wash, samples were dried for 20 min in a vacuum centrifuge (Vacufuge Concentrator 5301, Eppendorf, Hauppauge, NY) at 45°C. PCR Master Mix 1 (see online protocol) was added directly to the disks immediately prior to thermal cycling. Figure 1 shows that all tested cellulose matrices were an acceptable medium to capture, store, and amplify DNA. Disk washing was critical because unwashed disks had no detectable amplification products. Using water for the last two washes yielded the least consistent results, especially with untreated chromatography paper, while TE_0.1_yielded the best amplification regardless of matrix (Figure 1). To further streamline the process, disks from the same DNA sample can be washed together (one 1.5-mm disk to 100 μL of wash) to increase throughput.

**Figure 1.**
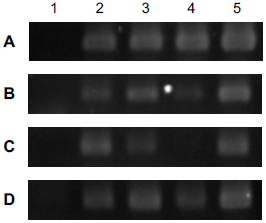
Comparison of four cellulose matrices and five washing conditions for isolation of DNA for PCR. *Arabidopsis thaliana* ecotype Columbia-0 leaf tissue was pressed onto (A) commercial FTA^®^ Card, (B) treated chromatography paper, (C) untreated chromatography paper, or (D) untreated filter paper. Initial wash treatments were: (1) unwashed; (2-5) washed twice with TENT buffer. Final wash treatments were: (1) unwashed; (2) isopropanol, (3) ethanol,(4) water or (5) TE_0.1_. PCR products were visualized on a 1.5% agarose gel using ethidium bromide.

Next, we compared the ability to store and amplify DNA from a variety of plant species on FTA^®^ Cards versus treated or untreated chromatography paper (Figure 2). All disks were washed twice with TENT buffer and twice with TE_0.1_. Two sequential PCRs were performed using pairs of nested primers to amplify a glyceraldehyde-3-phosphate dehydrogenase (*GAPDH*) gene (see online protocol). Many plants have more than one *GAPDH* gene (8-10), so PCR was first performed using purified genomic DNA as template to determine number and size of the expected products. The same products, ranging from 600 to 1600 bp depending upon the species, were detected from all matrices imprinted with *A. thaliana, P. sativum, C. sativus* or *C. quinoa* leaf tissue. In contrast, no product was generated from *L. sativa* or *O. basilicum* using any cellulose matrix. We noticed that the amount of moisture in the leaves of these two species was not sufficient to penetrate completely through the cellulose matrix. As good penetration is critical for optimum amplification, placing leaf tissue on both sides of the cellulose matrix before pressing can improve the results for less succulent leaves (11). For all remaining plants, the size and/or number of PCR products varied between matrices, so for these species the optimal matrix should be determined empirically.

**Figure 2.**
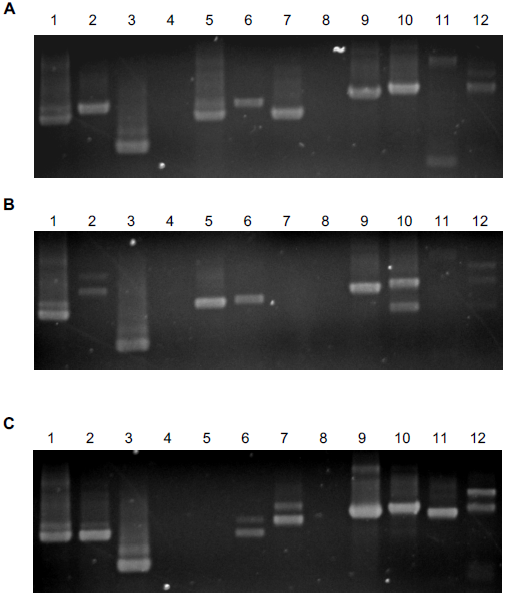
Comparison of three cellulose matrices for amplification of *GAPDH* genes from a variety of plants. Three different cellulose matrices were tested: (A) FTA^®^ Card, (B) treated chromatography paper, or (C) untreated chromatography paper. Young leaves from twelve different plants were imprinted on each cellulose matrix: (1) *Arabidopsis thaliana* ecotype Columbia-0, (2) *Physalis philadelphica* (tomatillo), (3) *Pisum sativum* var. *macrocarpon* cv. Oregon Sugar Pod (snow pea), (4) *Lactuca sativa* cv. Buttercrunch (lettuce), (5) *Avena sativa* (oats), (6) *Solanum lycopersicum* cv. Italian Heirloom (tomato), (7) *Medicago sativa* (alfalfa), (8) *Ocimum basilicum* cv. Genovese (basil), (9) *Cucumis sativus* cv. Suhyo Long (cucumber), (10) *Cucurbita pepo* cv. Early Yellow Crookneck (summer squash), (11) *Beta vulgaris* subsp. *vulgaris* cv. Ruby Red (Swiss chard), and (12) *Chenopodium quinoa* (quinoa). All disks were washed twice with TENT buffer and twice with TE_0.1_. PCR products were visualized on a 1.5% agarose gel using ethidium bromide.

Our results showed that: i) grade 238 chromatography paper pre-treated with Tris-EDTA-SDS buffer is a reasonable substitute for Whatman FTA^®^ Cards for many plant species, ii) homemade TENT buffer is a reliable replacement for FTA^®^ Purification Reagent for the first two disk washes, and iii) TE_0.1_ is a good choice for the last two washes. Sample processing time can be reduced by processing multiple disks from the same sample together and by using a vacuum centrifuge to dry the disks. A detailed protocol is available online.

## Acknowledgements

We thank the GEN606 class at the University of New Hampshire for amplifying the GAPDH gene from purified genomic DNA.

